# Exploring the Correlation between the Cognitive Benefits of Drug Combinations in a Clinical Alzheimer Disease Database and the Efficacies of the Same Drug Combinations Predicted from a Computational Model

**DOI:** 10.1101/437624

**Authors:** Thomas J. Anastasio

## Abstract

**INTRODUCTION:** Identification of drug combinations that could be effective in Alzheimer’s treatment is made difficult by the number of possible combinations. This analysis identifies as potentially therapeutic those drug combinations that rank highest when their efficacy is determined jointly from two independent data sources.

**METHODS:** Estimates of the efficacy of the same drug combinations were derived from a clinical dataset and from pre-clinical data, in the form of a computational model of neuroinflammation. Standard linear regression was used to show that the two sets of estimates were correlated, and to rule out possible confounds.

**RESULTS:** The ten highest ranking, jointly determined drug combinations most frequently consisted of COX2 inhibitors and aspirin, along with various antihypertensive medications.

**DISCUSSION:** Ten combinations of from five to nine drugs, and the three-drug combination of a COX2 inhibitor, aspirin, and a calcium-channel blocker, are discussed as candidates for consideration in future clinical and pre-clinical studies.

## 1. Introduction

Interest in polypharmacological approaches to the treatment of complex, multifactorial disorders is growing [eg 1]. Specific multi-target or multi-drug treatments for Alzheimer’s disease (AD) have already been suggested [2–6]. Combinations of approved, repurposed drugs could be more effective than single drugs in the treatment of AD, but determining which of the many possible combinations to use remains a challenge. The approach taken here is to combine clinical and pre-clinical data by correlating estimates of the epidemiological benefit of specific drug combinations, derived from a database, with predictions on their efficacy, derived from a computational model of a key component of AD pathophysiology.

Access to the database was provided by the Rush Alzheimer’s Disease Center (RADC database; https://www.radc.rush.edu/). RADC data was generated through the Religious Orders Study and Rush Memory and Aging Project [7]. The computational model was created on the basis of experimental data as published in the literature on microglia (MG model). Microglia mediate neuroinflammation, which is widely accepted as a key contributor to neurodegeneration and the resulting cognitive decline associated with AD [8–13]. Because estimates of drug combination benefit based on the RADC database are independent of the predictions of drug combination efficacy derived from the MG model, a positive correlation between the two would indicate drug combinations of potential value in treating AD. The analysis presented here integrates clinical and pre-clinical data and identifies novel combinations of widely prescribed drugs that stand as promising candidates for further pre-clinical or clinical evaluation.

## 2. Methods

### 2.1 Model structure and parameterization

The microglia model (MG model) is essentially a computational model of a cell, which can also be thought of as a model of many cells having exactly the same structure and function. Inputs impinging on the cell activates its receptors, which activate cell-signaling pathways, which activate transcription factors, which alter the cell’s expression of the proteins that the cell secretes and which can activate the cell’s own receptors, thus closing many positive and negative feedback loops. In the microglia model specifically, the secreted proteins are cytokines and other immunological factors that mediate the brain’s immune response and that can affect neurons and astrocytes but can also affect microglia themselves, via autocrine (one cell) or paracrine (many cells) feedback loops.

A highly simplified diagram of the microglia model is shown in Supplementary Fig. S1. The full microglia model is composed of 146 elements (units) that represent many of the receptors, signaling molecules, transcription factors, immunological factors (mainly cytokines), and some cellular processes (eg phagocytosis) that together determine the response of microglia to various inputs. A full list of model elements and abbreviations is provided in Supplementary Tab. S1. The model receives 90 inputs that represent endogenous or exogenous receptor ligands and also drugs and other compounds that bind receptors or that target other molecular entities (eg enzymes or transcription factors). All of the drugs and other compounds included in the model are listed in Supplementary Tab. S2.

The structure of the model incorporates the known interconnections between its elements that are described in the literature [for reviews see 14, 15–19], and it is an extension of previous models of microglia [20, 21]. The model takes the form of a recurrent network of nonlinear elements. The parameters of the model are the strengths (or weights) of the connections between model elements, and they are optimized by training the model using a machine-learning algorithm, which is specifically a recurrent neural network learning algorithm [22, 23]. A description of the parameter optimization procedure is provided in Supplementary Text S1.

The microglia model is trained using input/desired-output patterns that are derived from the results of *in vitro* (mainly) and *in vivo* experiments on microglia as described in the literature. A highly simplified input/desired-output table is shown in Supplementary Tab. S3. The full input/desired-output table has 179 entries. The connection weights are randomized prior to training, and the input/desired-output patters are presented many times and in random order during training (Supplementary Text S1). The goal in network construction and parameter optimization is to use available, pre-clinical data both to structure and to train the model so as to achieve the best possible representation of the function of microglia (Supplementary Text S2).

After training, the actual outputs of a trained network match the desired outputs with low error (Supplementary Fig. S2). Thus, a trained network can reproduce the known behavior of microglia as represented in the input/desired-output table, and can be used to predict the responses of microglia to novel inputs. Due to the randomness inherent in initial connection weight randomization, and to the random order of input/desired-output pattern presentations, predictions on the responses to novel inputs are improved when the responses of several trained networks are averaged [24]. The MG model results reported here are based on the averaged outputs of a set of ten networks, each trained from a different initial weight randomization and according to a different random schedule of input/desired-output presentations. To verify that this averaging procedure eliminated the bias inherent in predictions based on single networks, the results based on the averaged outputs of the set of ten networks that are reported here were compared with results based on the averaged outputs of a second set of ten networks, each trained using random initial weights and random input/desired-output presentation schedules that were different from each other and from those of the first set. The results derived from both sets of ten networks were highly consistent.

### 2.2 Predicting drug combination efficacy using the model

Any input to the microglia model can be thought of as a pattern over the values assigned to the 90 model input elements. Likewise, any output from the microglia model can be thought of as a pattern over the responses of the 18 units that are designated as model outputs (Supplementary Fig. S2). Two specific model output patterns are the neurotoxic and the neuroprotective patterns, which correspond to the responses of actual microglia that cause them to adopt, respectively, a highly pro-inflammatory or a highly anti-inflammatory response pattern.

Experimental results [25–27] and prior modeling [20, 21] suggest that the most potent pro-inflammatory stimulus, which may also represent actual conditions in the aging and AD brain [10, 28, 29], is a combination of amyloid-β (Aβ) and lipopolysaccharide (LPS), while the most potent anti-inflammatory stimulus is externally applied insulin-like growth factor-1 (IGF1). To assess the efficacy of any drug or drug combination in reducing the pro-inflammatory response, it is included in the input pattern along with the pro-inflammatory stimulus, Aβ and LPS, and the actual output is determined. The predicted efficacy of any drug or drug combination can then be defined as the amount by which it moves the response of the microglia model from the neurotoxic (highly pro-inflammatory) to the neuroprotective (highly anti-inflammatory) output pattern [21]. Specifically, the MG model efficacy of any drug combination is quantified as a ratio of normalized differences between the actual output response pattern and the neurotoxic and neuroprotective patterns, expressed as vectors (Supplementary Text S3 and Fig. S3). For reasons to be explained in the next subsection, a total of 196 drug combinations are included in the main analysis. The MG model efficacies of all 196 drug combinations included in the analysis range between 0.0682 and 0.5660, and the mean and variance are 0.3326 and 0.0151, respectively.

### 2.3 Assessing drug combination benefit from the database

The RADC dataset consists of up to 25 different assessments of the cognitive function of older participants, along with age (range 50 to 110 years), certain other demographic variables, a list of comorbidities, and self-reports of drug usage that were recorded on the initial visit and for some number of yearly follow-up visits thereafter [30, 31]. The 25 different cognitive function assessments, which had different measurement scales, were rescaled into the [0, 1] range and combined into a composite cognitive score for each visit (Supplementary Text S4).

The RADC dataset has nine comorbidity fields: hypertension, cancer (all types), diabetes, head injury, thyroid disease, congestive heart failure, claudication (peripheral vascular disease), heart disease (heart attack, myocardial infarction, etcetera), and stroke. Each binary comorbidity field contains a 1 if the participant self-reported that comorbidity and contains a 0 otherwise. The self-report indicated past history on the initial visit and any persisting or new comorbidity on subsequent visits. A simple comorbidity score was calculated by summing all the comorbidity fields for each visit.

The RADC dataset included data from 3326 participants whose drug usage was reported. In the RADC database the drugs participants reported taking were grouped into 100 drug categories, many of which were redundant or otherwise overlapping. Of those 100 drug categories, 20 were chosen because they were relatively non-overlapping, and because the effects on microglia of one or more of the drugs from that category had been determined. The 20 chosen drug categories, with names close to those used in the RADC database, are: acetaminophen, COX2 inhibitors, antimalarials, aspirin, glucocorticoids, opioids, antibiotics, ACE inhibitors, anti-adrenergics, beta blockers, calcium-channel blockers, angiotensin-receptor blockers, anti-arrhythmics, anti-diabetics, estrogen, spironolactone, proton-pump inhibitors, antimanics, antidepressants, and antihistamines. Any overlaps between these 20 drug categories were considered admissible because they would be expected to weaken rather than falsely strengthen any agreement between RADC database benefits and MG model efficacies, and because they were relatively minor overall.

The RADC database recorded the drugs in each category that the participant reported using on each visit in a binary fashion (1 if taken, 0 if not). To get a composite view of the drugs used by each participant, the drug usage over all visits were combined using a logical OR, so that a participant was designated as a user of a drug of a specific class if that participant had reported using a drug from that class on at least one visit. The simplifying assumption here is that the effect of any drug on cognitive function (or on neurodegeneration or neuroinflammation) is independent of the duration of use of that drug. This assumption is almost certainly false but was considered admissible because it would weaken rather than falsely strengthen any agreement between RADC database benefits and MG model efficacies.

When drug combination usage was combined in this way, using the logical OR over all visits, 2167 of the possible 1,048,576 combinations of the 20 drug types were actually used by RADC participants. For simplicity of exposition, the term “drug combination” will subsume combinations of two or more drugs as well as single drugs. RADC participants were grouped according to drug combination, yielding 2167 different drug combination groups containing at least one participant whose age and composite cognitive score were recorded on at least one visit.

The benefit of any drug or drug combination to RADC participants was assessed as the difference in cognitive function of participants who reported taking that drug or drug combination, and the cognitive function of participants who reported taking no drugs. The well-known decline of cognitive function with age [30, 31] for all of the participants in each drug combination group was summarized by pooling all of the composite cognitive score versus age (cog-score vs age) values for all participants in each drug combination group, and fitting them with a simple, three-parameter power function via nonlinear regression (Supplementary Text S4 and Fig. S4). Because at least three data points are needed to specify a three-parameter power function, any drug combination having fewer than three cog-score vs age values was removed, leaving 1955 drug combination groups (including the no-drug group). As a further safeguard, only dug combination groups containing at least three participants were given further consideration, leaving a total of 196 drug combination groups in the main analysis.

The power function curve for each drug combination group was used to compute the expected cog-score at each age included in that specific drug combination group. Also, the power function curve for the no-drug group was used to the compute the expected cog-score in the no-drug case, but at the same ages as were included for that specific drug combination group. Then the expected cog-score in the no-drug case was subtracted from the expected cog-score for that specific drug combination *at the same set of ages*, and the differences were averaged to yield the RADC database benefit for that drug group. Note that many of the RADC database benefits are negative when computed this way, so the relative rather than the absolute values of the RADC database benefits are relevant to this analysis. For the 196 drug combinations that were included in the main analysis, RADC database benefits range from −0.4541 to 0.1173, and the mean and variance are −0.0260 and 0.0063, respectively.

All computations were performed using MATLAB™ version R2017b, running on an Intel i5-7500T quad-core CPU at 2.7 GHz per core with 16 GB RAM.

## 3. Results

### 3.1 Correlating MG model efficacies and RADC database benefits

MG model efficacies and RADC database benefits are significantly correlated. Fig. 1 shows the statistically significant correlation for all drug combinations taken by three or more RADC participants and having three or more cog-score vs age values, pooled over the participants in the corresponding drug combination group. Each data point in Fig. 1 corresponds to a unique drug combination, and is located in the plot according to its MG model efficacy and its RADC database benefit. The slope of the regression line (s) is 0.1339, the degree of linearity (or correlation coefficient, r) is 0.2064, and the probability that the correlation occurred by random chance (p) is 0.0037. The set of 196 data points on which this correlation is based will henceforth be called the “correlation set”.

As explained above, several assumptions are implied in making a comparison between MG model efficacy and RADC model benefit, but most of them would be expected to degrade the correlation. The salient exception would be a spurious correlation related to the fact that, due to experimental bias, most of the drugs included in the MG model are those that have been shown to have anti-inflammatory effects on microglia, which are considered positive results in the literature. The bias against reporting negative results means that data on drugs that have a pro-inflammatory effect on microglia are relatively very few.

**Fig. 1.**
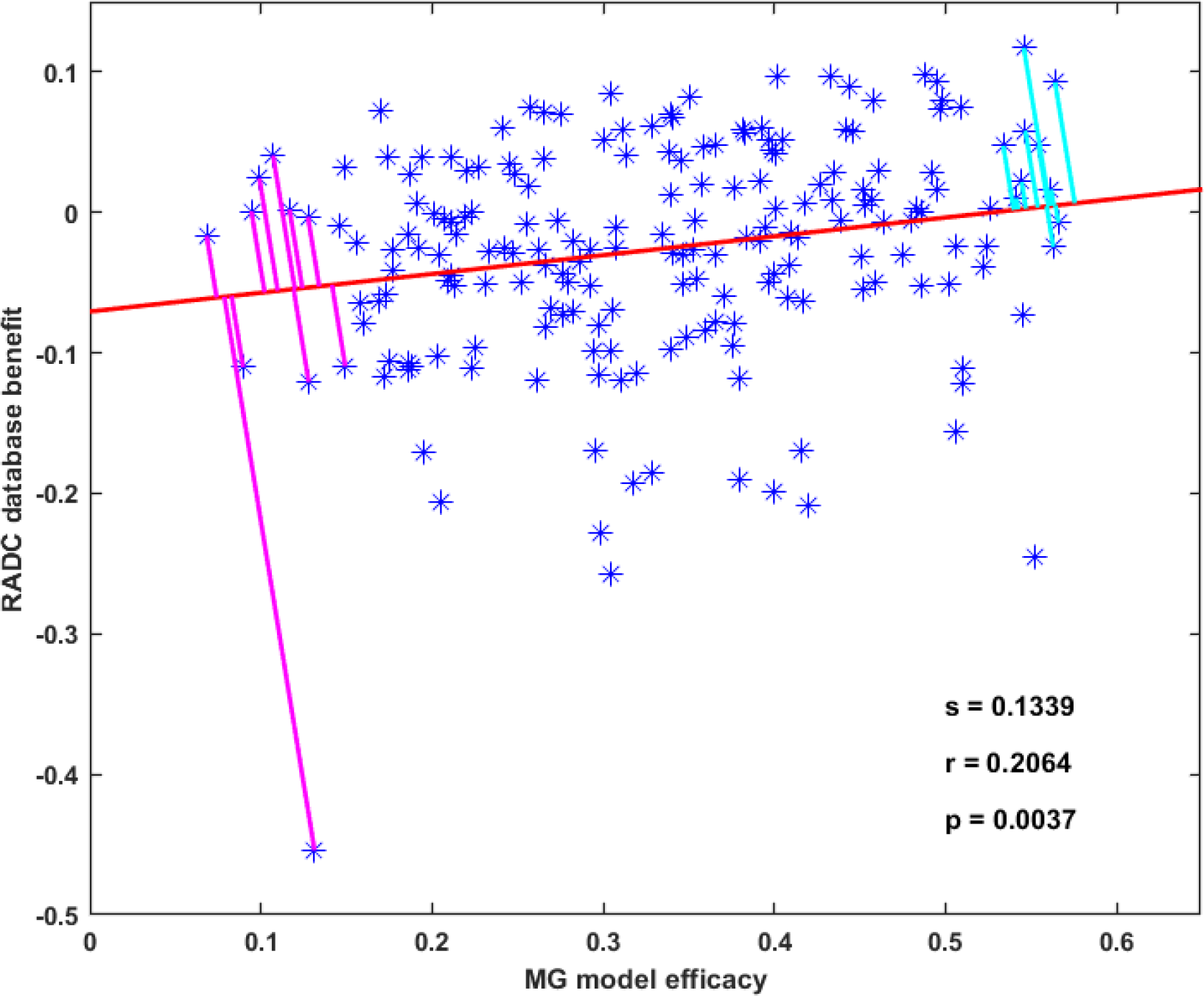
MG model efficacies and RADC database benefits are significantly correlated. Each point represents one of the 196 drug combinations that was included in the main analysis. The regression line fit to the data points has slope s = 0.1339, correlation coefficient r = 0.2064, and p-value p = 0.0037. The line segments perpendicular to the regression line shown the Ten Best (upper right) and the Ten Worst (lower left) drug combinations as determined jointly by the MG model and the RADC database.

As noted above, only 2167 of the possible 1,048,576 combinations of drugs in the 20 categories included in the analysis were actually taken by RADC participants, and of those only 196 combinations were taken by at least three participants and included at least three cog-score vs age data points. A computational screen using the MG model over all combinations of the drugs in common between the model and the RADC dataset shows that the model may indeed capture potential antagonisms between the drugs, such that the anti-inflammatory effect of single drugs may actually be reduced in some combinations (Supplementary Fig. S5). However, over the much more limited range of the 196 drug combinations in the correlation set, which has a maximal number of drugs per combination of ten, the efficacies of drug combinations in the MG model tend to rise as the number of drugs in the combination rises. This is shown in Fig. 2A. A spurious correlation between MG model efficacies and RADC database benefits could result if RADC benefit also tended to rise as the number of drugs in the combination rises. However, the RADC benefits of the drug combinations in the correlation set stay constant as the number of drugs in the combination rises. This is shown in Fig. 2B.

**Fig. 2.**
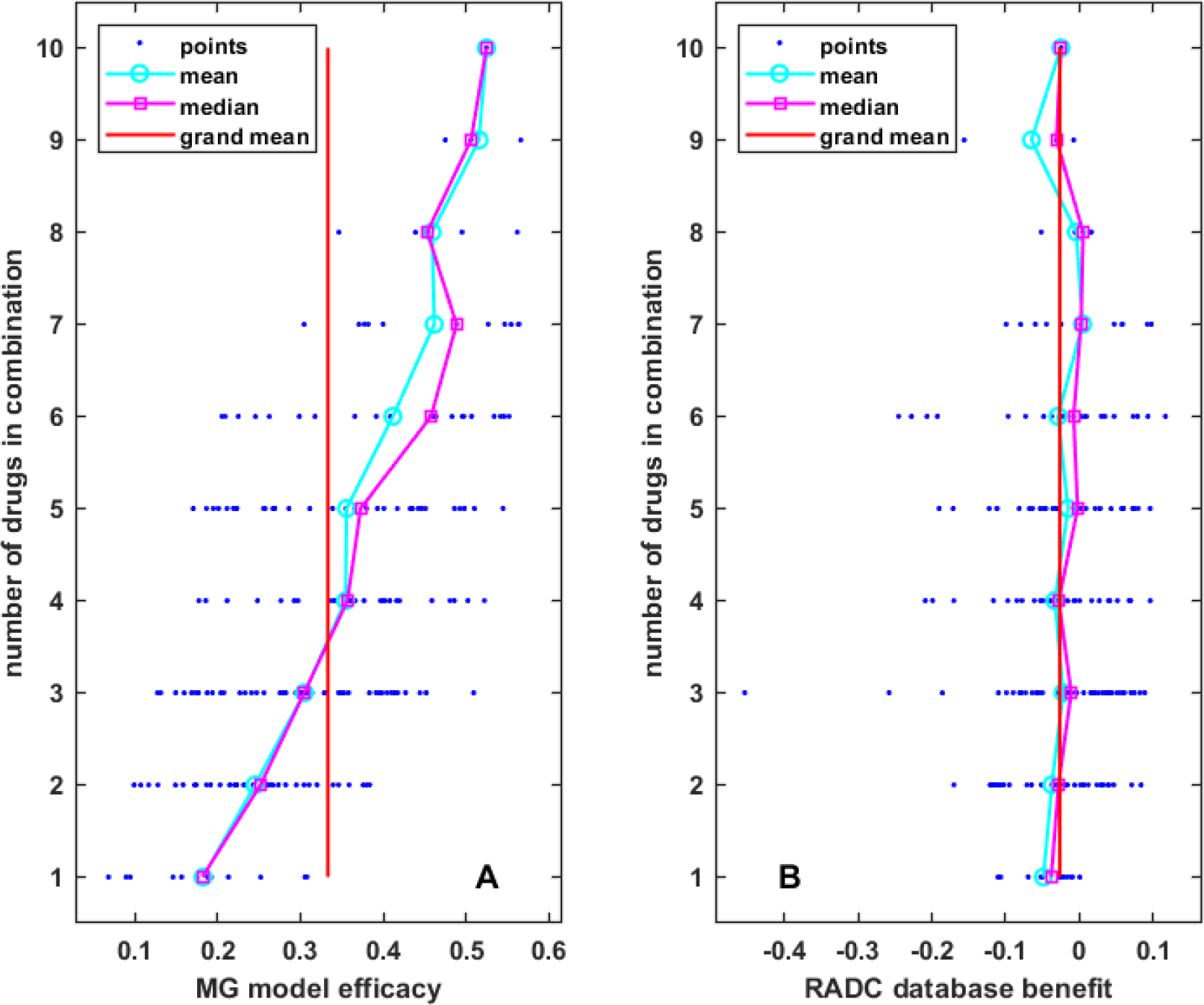
MG model efficacy and RADC database benefit vary widely for drug combinations that are composed of the same number of drugs. In both panels, circles and squares are the mean and median over drug combinations composed of the same number of drugs, and the vertical line is the grand mean over all drug combinations. Mean MG model efficacy rises (A) but mean RADC database benefit stays constant (B) as the number of drugs in the drug combinations rises.

The lack of correlation between clinically observed benefit and number of drugs in the combinations included in the correlation set reflects a lack of correlation between clinically observed cognitive function and number of drugs over the whole RADC dataset. The results of some regression analyses of the whole RADC dataset is shown in Fig. 3. They are based on the 8675 entries in the RADC dataset for which complete information on age, cognition, comorbidities, and drugs taken is available. There was a strong and highly statistically significant correlation (s = 2.0454, r = 0.3697, p = 0.0000) between the number of drugs taken by a participant and their combined comorbidity score, as shown in Fig. 3A. This should be expected for the RADC dataset because it is a sample of a North American population that generally has good access to medical care and to prescription drugs. The correlation between the number of drugs taken by a participant and their age was also highly statistically significant but was weak (s = 0.0843, r = 0.0868, p = 0.0000), as shown in Fig. 3B.

**Fig. 3.**
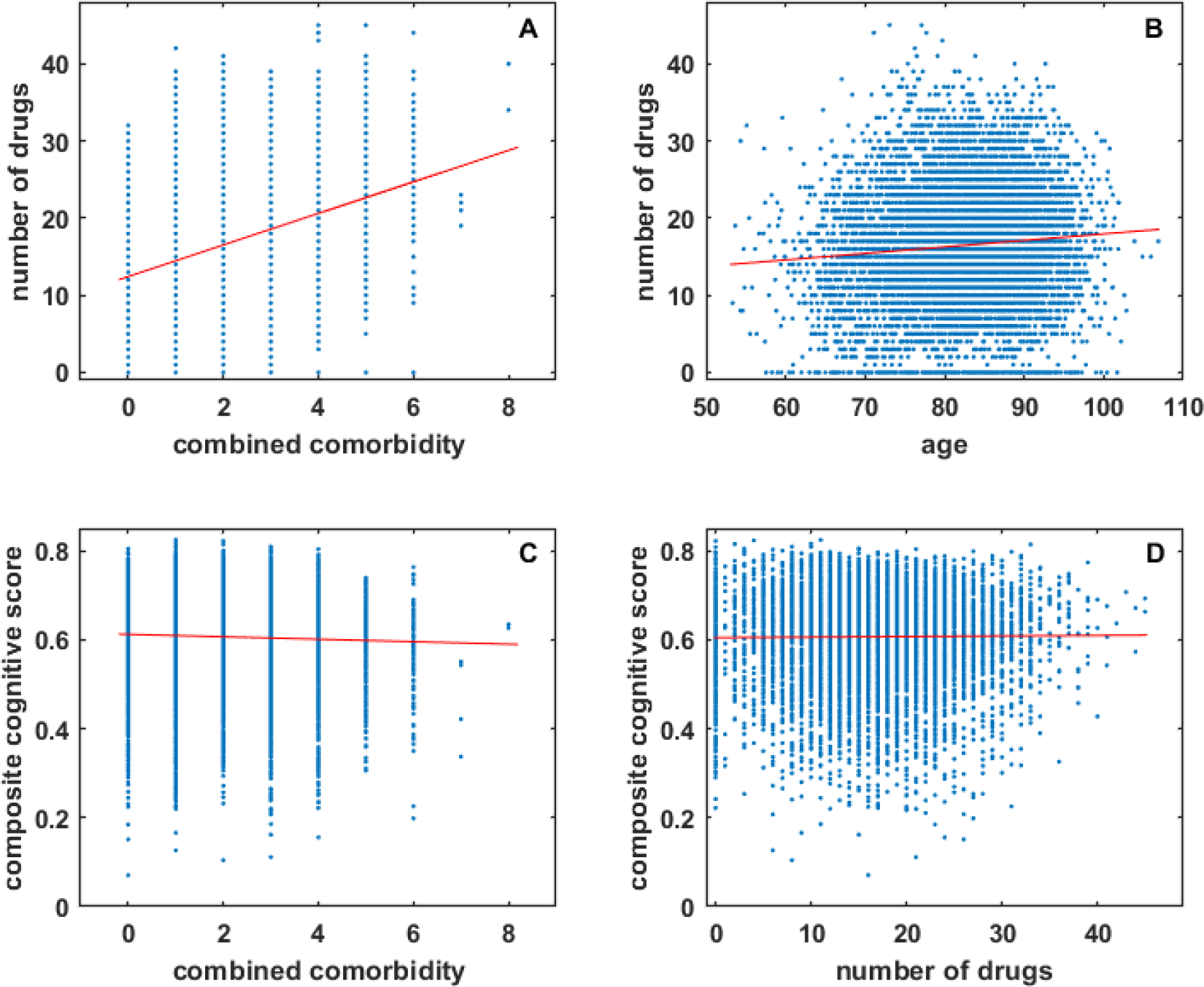
RADC participants took more drugs as they aged and developed comorbidity, but their cognitive scores were not strongly dependent on comorbidity or number of drugs taken. The number of drugs taken by RADC participants overall increases with comorbidity (A) or with age (B), but composite cognitive score is only very weakly correlated with comorbidity (C), and there is no correlation between composite cognitive score and the number of drugs taken by a RADC participant (D).

Analysis of the whole RADC dataset suggests that cognitive performance degrades with comorbidity, as shown in Fig. 3C. Although the correlation is statistically significant it is very weak (s = −0.0028, r = −0.0363, p = 0.0007; see also below). Of central concern here is any relationship between cognition and the number of drugs taken, but regression analysis clearly indicates (s = 0.0001, r = 0.0091, p = 0.3957) that there is no such relationship for the RADC dataset, as shown in Fig. 3D. The very weak relationship between cognition and comorbidity, and the lack of relationship between cognition and the number of drugs taken, justifies the pooling of RADC participants over comorbidities and over drugs taken outside of the 20 categories included in this analysis.

The same lack of relationship between cognition and the number of drugs taken is shown in Fig. 2B for the correlation set, where it is also apparent that both RADC database benefit and MG model efficacy vary over a broad range for drug combinations composed of the same number of drugs. These graphical and regression analyses show that it is not simply the number of drugs in the combinations, but the specific drugs in the combinations, which jointly determine their MG efficacy and RADC benefit.

### 3.2 Finding the Ten Best and Ten Worst drug combinations

Having established a statistically significant correlation between MG model efficacy and RADC database benefit, the regression line can be used to estimate the efficacy of each drug combination using both measures together rather than either one separately. This joint efficacy measure can be determined by finding the distances from the y-intercept of the projection of each drug combination data point onto the regression line. Then the Ten Best and Ten Worst drug combinations, determined jointly according to MG efficacy and RADC benefit, are those whose projections have the longest and shortest distances, respectively, along the regression line (Fig. 1). The drugs that compose the Ten Best and Ten Worst drug combinations are indicated in Tab. 1.

**Tab. 1.** The Ten Best and Ten Worst drug combinations as determined jointly from MG model efficacies and RADC database benefits. When determined jointly, the Ten Best drug combinations are composed of more drugs than the Ten Worst combinations. The presence of COX2 inhibitors, aspirin, and calcium-channel blockers saliently distinguishes the Ten Best drug combinations. The presence of opioids distinguishes the Ten Worst. The drug category names are close to those assigned in the RADC database, and are followed in parentheses by the drug (or drugs) that represent that category in the MG model.

| rank | acetaminophen (acetaminophen) | COX2 inhibitor (ibuprofen) | antimalarial (chloroquine) | aspirin (aspirin) | glucocorticoid (dexamethasone) | opioid (morphine) | antibiotic (minocycline/rifampicin) | ACE inhibitor (perindopril) | anti-adrenergic (clonidine) | beta blocker (propranolol) | calcium-channel blocker (nifedipine) | angiotensin-receptor blocker (losartan) | anti-arrhythmic (flecainide) | antidiabetic (rosiglitazone/glimperide) | estrogen (estrogen) | spironolactone (spironolactone) | proton-pump inhibitor (omeprazole) | antimanic (lithium) | antidepressant (fluoxetine) | antihistamine (diphenhydramine) |
| --- | --- | --- | --- | --- | --- | --- | --- | --- | --- | --- | --- | --- | --- | --- | --- | --- | --- | --- | --- | --- |
| **TEN BEST DRUG COMBINATIONS DETERMINED JOINTLY (MG AND RADC)** |  |  |  |  |  |  |  |  |  |  |  |  |  |  |  |  |  |  |  |  |
| 1 | X | X |  | X |  |  |  | X |  | X | X | X |  |  |  |  |  |  |  |  |
| 2 | X | X |  | X |  |  | X | X | X | X | X |  |  |  |  |  |  |  |  | X |
| 3 | X | X |  | X |  |  |  |  |  | X | X | X |  |  |  |  | X |  | X |  |
| 4 |  | X |  | X |  |  |  |  |  | X | X | X |  |  |  |  | X |  |  |  |
| 5 | X | X |  | X |  |  |  |  |  | X | X | X |  |  |  |  |  |  | X |  |
| 6 | X | X |  | X |  |  |  | X |  | X | X |  |  | X |  |  |  |  |  |  |
| 7 | X | X |  | X |  |  | X | X |  | X | X |  |  |  |  |  |  |  |  |  |
| 8 | X |  |  | X |  |  |  | X |  |  | X |  |  | X |  |  |  |  |  |  |
| 9 |  |  |  | X |  |  |  | X |  | X | X |  |  | X |  |  | X |  |  |  |
| 10 | X |  |  | X |  |  | X |  |  | X | X | X |  |  |  |  |  |  |  |  |
| **TEN WORST DRUG COMBINATIONS DETERMINED JOINTLY (MG AND RADC)** |  |  |  |  |  |  |  |  |  |  |  |  |  |  |  |  |  |  |  |  |
| 1 |  |  |  |  |  |  |  |  |  | X |  |  |  |  |  |  |  |  |  |  |
| 2 | X |  |  |  |  | X |  |  |  |  |  |  |  |  |  |  |  |  | X |  |
| 3 | X |  |  |  |  |  |  |  |  |  |  |  |  |  |  |  |  |  |  |  |
| 4 |  |  |  |  |  |  |  |  |  |  |  |  |  |  |  |  | X |  |  |  |
| 5 | X |  |  |  |  |  |  |  |  | X |  |  |  |  |  |  |  |  |  |  |
| 6 | X |  |  |  |  |  |  |  |  |  |  |  |  |  |  |  | X |  |  |  |
| 7 |  |  |  |  |  |  |  |  |  | X |  |  |  |  |  |  | X |  |  |  |
| 8 |  |  |  |  |  |  |  | X |  | X |  |  |  |  |  |  |  |  |  |  |
| 9 | X |  |  |  |  |  |  |  |  | X |  |  |  |  |  |  | X |  |  |  |
| 10 | X |  |  | X |  | X |  |  |  |  |  |  |  |  |  |  |  |  |  |  |

The number of drugs in the Ten Best combinations have a mean of seven and range from five to nine, while that for the Ten Worst have a mean of two and range from one to three. The percentages of the drugs in the Ten Best and Ten Worst drug combinations are shown relative to each other and to the overall drug percentages in Fig. 4. In Fig. 4 as in Tab. 1, COX2 inhibitors, aspirin, and calcium-channel blockers stand out because they are represented at higher percentages in the Ten Best than overall and much higher than in the Ten Worst. Opioids are represented at a lower percentage in the Ten Worst drug combinations than overall but are absent from the Ten Best combinations.

**Fig. 4.**
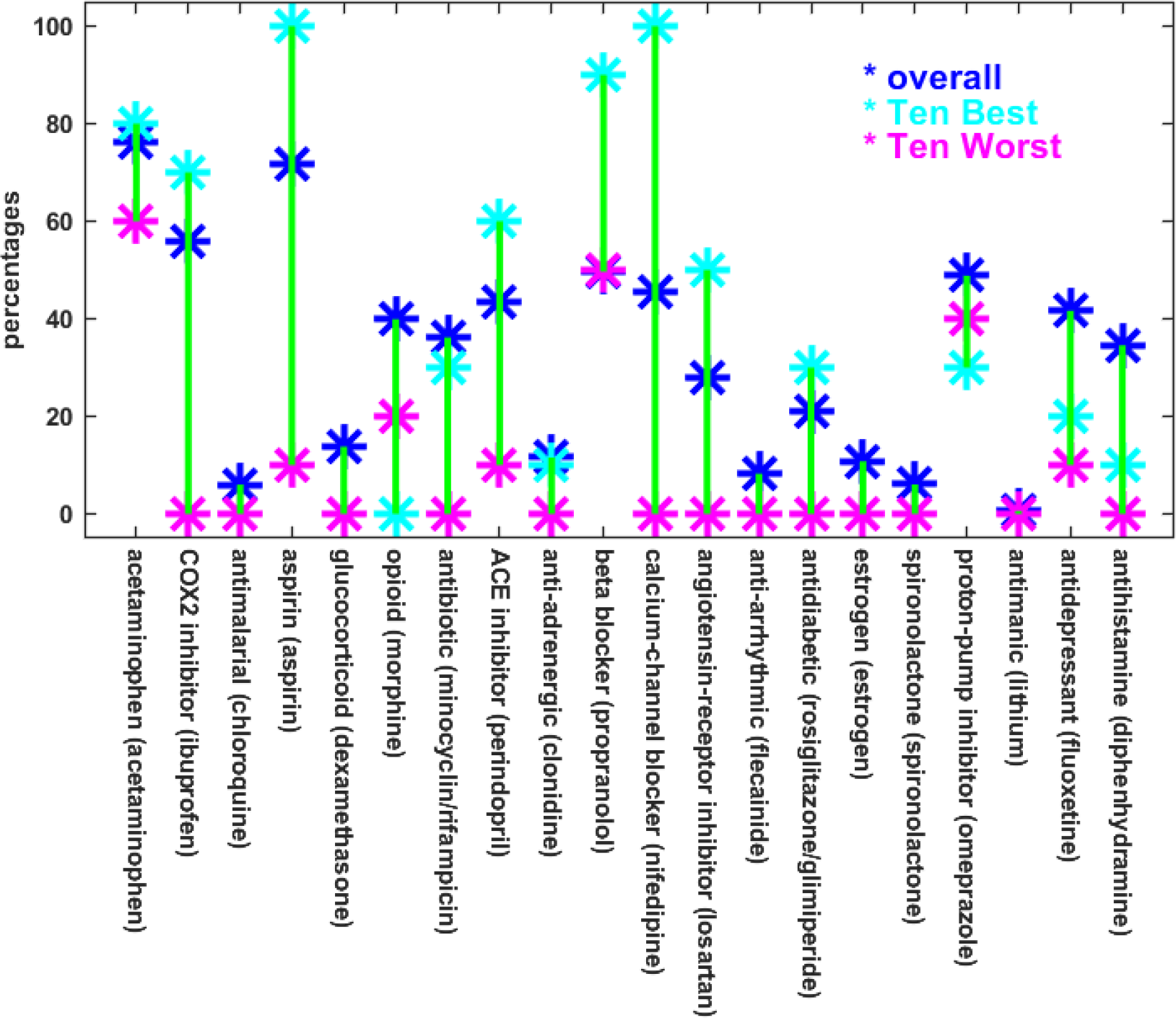
The percentages of drugs that appear in the Ten Best and Ten Worst combinations are different from their overall percentages. This is especially true for aspirin and calcium-channel blockers, which appear at much higher percentages than overall in the Ten Best combinations and at much lower percentages than overall in the Ten Worst combinations. A similar but less pronounced relationship is observed for COX2 inhibitors, ACE inhibitors, and angiotensin-receptor blockers. The reverse relationship holds for the opioids. The labels along the bottom are similar to the RADC drug category designations, followed by the name of the specific drug included in the MG model to represent that category. Note that the MG model included both minocycline and rifampicin for the antibiotic RADC category, and both rosiglitazone and glimepiride for the antidiabetic category, and the averaged outputs for each pair of drugs represented the response of the model for the corresponding category.

The regression analysis in Fig. 1 suggests that the Ten Best and Ten Worst drug combinations as determined jointly from MG model efficacy and RADC database benefit would be more similar to the ten best if determined from the MG model alone than they would be if determined from the RADC dataset alone. Indeed, if the ten best are determined from either the MG model or RADC dataset alone, then the Ten Best as determined jointly has eight of the ten best MG model efficacies but only two of the ten best RADC database benefits. Similarly, if the ten worst are determined from either the MG model or RADC database alone, then the Ten Worst as determined jointly has nine of the ten worst MG model efficacies but only one of the ten worst RADC database benefits.

The ten best and ten worst drug combinations as determined from the RADC database alone are indicated in Tab. 2. Some differences are apparent. Whereas the Ten Best drug combinations are composed of more drugs than the Ten Worst, when determined jointly from the MG model and RADC database (mean Best is seven, mean Worst is two), the ten best and ten worst have about the same number of drugs when determined from the RADC database alone (mean best and mean worst are both five). Also, whereas acetaminophen and antidepressants are more frequent in the Ten Best than in the Ten Worst jointly determined drug combinations, they are less frequent in the ten best than in the ten worst RADC-alone determined drug combinations, and the best/worst differences in antidepressants is especially dramatic for the RADC-alone determined combinations.

**Tab. 2.** The ten best and ten worst drug combinations determined from RADC database benefits only. The ten best and ten worst RADC-alone determined drug combinations are composed of similar numbers of drugs. Calcium-channel blockers occur more frequently in the ten best (5/10) than in the ten worst (1/10). Opioids are notably absent from the ten best but present in almost half of the ten worst, and antidepressants are absent from the ten best but frequent in the ten worst. The drug category names are close to those assigned in the RADC database, and are followed in parentheses by the drug (or drugs) that represent that category in the MG model.

| rank | acetaminophen (acetaminophen) | COX2 inhibitor (ibuprofen) | antimalarial (chloroquine) | aspirin (aspirin) | glucocorticoid (dexamethasone) | opioid (morphine) | antibiotic (minocycline/rifampicin) | ACE inhibitor (perindopril) | anti-adrenergic (clonidine) | beta blocker (propranolol) | calcium-channel blocker (nifedipine) | angiotensin-receptor blocker (losartan) | anti-arrhythmic (flecainide) | antidiabetic (rosiglitazone/glimiperide) | estrogen (estrogen) | spironolactone (spironolactone) | proton-pump inhibitor (omeprazole) | antimanic (lithium) | antidepressant (fluoxetine) | antihistamine (diphenhydramine) |
| --- | --- | --- | --- | --- | --- | --- | --- | --- | --- | --- | --- | --- | --- | --- | --- | --- | --- | --- | --- | --- |
| **TEN BEST DRUG COMBINATIONS DETERMINED FROM RADC ALONE** |  |  |  |  |  |  |  |  |  |  |  |  |  |  |  |  |  |  |  |  |
| 1 |  | X |  | X |  |  |  | X |  | X | X |  |  |  |  |  |  |  |  |  |
| 2 |  |  |  | X |  |  |  |  |  | X |  | X |  |  |  |  |  |  |  |  |
| 3 |  |  |  | X |  |  | X |  |  |  |  |  |  |  |  |  |  |  |  |  |
| 4 |  | X |  | X |  |  |  |  |  |  | X |  |  |  |  |  |  |  |  |  |
| 5 | X | X |  | X |  |  |  | X |  | X | X | X |  |  |  |  |  |  |  |  |
| 6 | X | X |  | X |  |  |  | X |  | X |  |  |  | X |  |  |  |  |  |  |
| 7 | X |  |  | X |  |  |  | X |  | X | X |  |  |  |  |  |  |  |  |  |
| 8 |  | X |  | X |  |  |  |  |  |  |  |  |  |  |  |  | X |  |  | X |
| 9 | X | X |  | X |  |  | X | X |  | X |  |  |  |  |  |  | X |  |  |  |
| 10 |  | X |  | X |  |  |  |  |  | X | X | X |  |  |  |  | X |  |  |  |
| **TEN WORST DRUG COMBINATIONS DETERMINED FROM RADC ALONE** |  |  |  |  |  |  |  |  |  |  |  |  |  |  |  |  |  |  |  |  |
| 1 | X |  |  |  |  | X |  |  |  |  |  |  |  |  |  |  |  |  | X |  |
| 2 | X |  |  | X |  |  |  |  |  |  |  |  |  |  |  |  |  |  | X |  |
| 3 | X | X |  | X |  |  |  | X |  |  | X |  |  |  |  |  |  |  | X |  |
| 4 | X | X |  | X |  | X |  |  |  |  |  |  |  |  |  |  | X |  | X |  |
| 5 |  |  |  | X |  |  |  | X |  | X |  |  |  | X |  |  |  |  |  |  |
| 6 | X |  |  | X |  | X |  |  |  | X |  |  |  |  |  |  | X |  | X |  |
| 7 | X | X |  | X |  |  |  |  |  |  |  |  |  |  |  |  |  |  | X |  |
| 8 | X |  |  | X |  | X | X |  |  |  |  |  |  |  |  |  | X |  | X |  |
| 9 | X | X |  | X |  |  |  |  |  | X |  |  |  |  |  |  |  |  | X |  |
| 10 | X | X |  |  |  |  |  |  |  |  |  |  |  |  |  |  |  |  | X |  |

Many similarities between the jointly determined and RADC-alone determined estimates are also apparent. COX2 inhibitors, aspirin, antibiotics, ACE-inhibitors, beta blockers, calcium-channel blockers, angiotensin receptor inhibitors, and antihistamines, which are all present at higher frequencies in the Tens Best than in the Ten Worst jointly determined drug combinations, are also all present at higher frequencies in the ten best than in the ten worst RADC-alone determined drug combinations. The opioids, which are present at lower frequency in the Ten Best than in the Ten Worst jointly determined drug combinations, are also present at lower frequency in the ten best than in the ten worst RADC-alone determined drug combinations.

### 3.3 Ruling out hypertension and other possible confounds

The Ten Best jointly determined (MG and RADC) and the ten best RADC-alone determined drug combinations are similar in that they include ACE-inhibitors, beta blockers, calcium-channel blockers, and angiotensin receptor inhibitors. While some studies disagree [32, 33], most studies find that these antihypertensive drugs considered singly have been associated with a lower risk of AD, and in some cases lower risk was independent of hypertension [34–39]. These drugs are frequently taken in combination to treat hypertension, and the question arises as to the potential influence of hypertensive status on the efficacy of the various drug combinations determined in this analysis.

Hypertension, like overall comorbidity, degrades cognitive performance but the effect is small. As noted above, the negative correlation between cognitive performance and comorbidity in the larger RADC dataset is significant (p = 0.0007) but very weak (s = −0.0028, r = −0.0363). Similarly, a negative effect on cognitive performance of hypertension is also observed in the larger RADC dataset, and the difference between the mean cognitive scores of participants with hypertension (0.6036) and without hypertension (0.6109) is significant (p = 0.0011) but very small. In contrast, there are strong, positive correlations between comorbidity (s = 1.2821, r = 0.3611, p = 0.0000), or hypertension specifically (s = 2.1258, r = 0.4551, p = 0.0000), and the efficacy of the drug combinations in the correlation set as determined jointly from the MG model and the RADC database (Supplementary Fig. S6 and Tab. S6).

The strong, positive correlations between comorbidity or hypertension and drug combination efficacy seem to imply that the groups of participants who take effective drug combinations have high cognitive scores because they also have high comorbidities or have a high proportion of hypertensives, but these implications are false because, as we have seen, comorbidity and hypertension are both negatively related to cognitive performance in the RADC dataset. In any case, the effects of comorbidity or hypertension on cognitive performance are very weak or very small and can be ignored in the analysis. The strong, positive correlations between comorbidity or hypertension and drug combination efficacy imply instead that the participants who suffer comorbidities generally, or hypertension superficially, tend to be those who take combinations of the drugs examined in this analysis that are known to reduce the pro-inflammatory responses of microglia. Further analysis of the demographic variables available in the RADC dataset rule out other possible confounds (Supplementary Fig. S6 and Tabs. S6-S8).

### 3.4 Focusing on specific drug combinations

Featured prominently in the Ten Best jointly determined, and in the ten best RADC-alone determined drug combinations are four antihypertensive drug types: ACE inhibitors, beta blockers, calcium-channel blockers, and angiotensin-receptor blockers. Unfortunately, none of the 196 combinations included in the correlation set comprised only those four drug types, with or without COX2 inhibitors and/or aspirin. The three single drugs that stand out in the best combinations, whether determined jointly or by RADC alone, are the COX2 inhibitors, aspirin, and calcium-channel blockers, while the drugs that stand out in the worst combinations are the opioids and the antidepressants, and the question arises as to whether or not combinations of just those few drugs would be especially good or bad by themselves. None of the three possible combinations of the opioids and the antidepressants (each alone and the one pair) is present in the correlation set. However, all seven possible combinations of the COX2 inhibitors, aspirin, and calcium-channel blockers (each alone, all three pairs, and the one triple) are present in the correlation set. Their projections (not shown) onto the regression line in Fig. 1 are distributed over the length of the line but none rise to the level of the Ten Best nor fall to the level of the Ten Worst.

The projections of each of the COX2 inhibitors, aspirin, and calcium-channel blockers by themselves (0.1436, 0.1870, and 0.2431) fall below those of each pair (0.2700, 0.3390, and 0.3606). The best of the seven possible combinations of the anti-inflammatory drugs, aspirin, and calcium-channel blockers is the triple composed of all three, and its projection (0.4521) is closest to those of the Ten Best (range 0.5353 to 0.5713). These findings imply that the triple of COX2 inhibitors, aspirin, and calcium-channel blockers would be better than those same drugs alone or in pairs as combination therapies for AD. They also imply that any of the Ten Best combinations, all of which include drugs in addition to COX2 inhibitors, aspirin, and calcium-channel blockers, would be better than the combination limited to the triple of COX2 inhibitors, aspirin, and calcium-channel blockers as combination therapies for AD.

### 3.5 Analysis of variance and multiple comparisons

The main challenge in the identification of potential multi-drug treatments for AD using clinical data is that the participants in the dataset are distributed over a great many drug combinations. The result is that the amounts of data associated with each individual drug combination is small, and this reduces statistical power. The problem is illustrated using the statistical analysis presented in Supplementary Figs. S7-S10 and Tabs. S7-S10. Fig. S9 shows that the medians of the combined cognitive scores of the ten best and ten worst RADC-alone drug combinations are all, respectively, higher and lower than the median for the “other” group (including other combinations and the no-drug case), and this agreement corroborates the method used to assess RADC dataset benefit (see Methods). However, using a multiple comparisons test with the Bonferroni correction shows that only two of the ten best are statistically significantly better than the other category (Tab. S9). As expected from Fig. 1, the relationships between the medians of the combined cognitive scores of the Ten Best and Ten Worst jointly determined drug combinations and that for the other group is not as clean (Fig. S10), but here again two of the Ten Best are statistically significantly better than the other category (Tab. S10). A multi-way analysis of variance also confirms the presence of interactions that were identified, and ruled out, above.

Despite lack of significance in this more traditional form of analysis, the relative differences in statistics such as medians and means (Supplementary Figs. S7-S10 and Tabs. S7-S10) supports the contention that certain drug combinations, particularly those including COX2 inhibitors, aspirin, and antihypertensive drug in categories known as ACE inhibitors, beta blockers, calcium-channel blockers, and angiotensin-receptor blockers, may be more effective than single drugs in the treatment of AD.

## 4. Discussion

The goal of this analysis was to take estimates of the efficacy/benefit of specific drug combinations as determined completely independently, using the MG model and the RADC database, and reduce the uncertainty inherent in either by correlating them both together. The main assumption in this analysis is that certain drug combinations as identified by the MG model can reduce neuroinflammation, which in turn can reduce neurodegeneration, which in turn can improve cognitive performance, and that the relative improvement in cognitive score should be detectable for those same drug combinations in the RADC dataset. The analysis of available data supports the main assumption, which is that the agreement between the MG model and the RADC database indicates that specific drug combinations are associated with relative cognitive benefit because they reduce neuroinflammation. The analysis does not rule out the possibility that certain drugs provide a relative benefit because they also ameliorate other aspect(s) of AD-related pathology beside neuroinflammation, as has been suggested for certain calcium-channel blockers [40, 41] and COX2 inhibitors [42–45].

COX2 inhibitors, aspirin, antibiotics, ACE-inhibitors, beta blockers, calcium-channel blockers, angiotensin receptor inhibitors, and antihistamines were all present at higher frequencies in the Ten Best than in the Ten Worst jointly determined drug combinations, and were also all present at higher frequencies in the ten best than in the ten worst RADC-alone determined drug combinations. All of these drugs have known anti-inflammatory effects on microglia and, while some have overlapping sets of targets, all have distinct targets, suggesting that their effects may synergize because they affect different cellular pathways (Supplementary Tab. S2 and references therein).

The mean number of drugs in the Ten Best drug combinations is seven, with a range of five to nine, and the mean number of drugs, limited to the 20 categories examined here, that were taken by RADC participants is five, and range up to 16, so prescribing and taking a combination of from five to nine drugs for the treatment of AD would not be inconsistent with current clinical practice in North America. However, the analysis also suggest that taking a combination of three drugs, namely COX2 inhibitors, aspirin, and calcium-channel blockers, would be more beneficial than taking any of those drugs alone.

High-throughput experiments on microglia, or on mixed neural/glial cultures, would be the most economical way to test the actual efficacy of the drug combinations identified in this analysis. For example, all combinations of a COX2 inhibitor, aspirin, and a calcium-channel blocker, with each drug at one of seven concentrations, could be tested in quadruplicate for statistical power on a single, 1536 microtiter plate. New clinical studies are also feasible. All of the Ten Best drug combinations include at least two of the following drugs: ACE inhibitors, beta blockers, calcium-channel blockers, and angiotensin-receptor blockers. These drugs are frequently taken in combination to treat hypertension, and many of the elderly patients likely to enter new clinical studies are already taking two or more of them together, along with some of the other drugs included in the Ten Best list. Adding drugs such as COX2 inhibitors and aspirin to already established drug regimens, in order to complete some of the Ten Best combinations, is an option for future clinical trials.

## 5. Acknowledgements

This work was supported by The Rotary Coins for Alzheimer’s Research Trust Fund, administered by the Rotarians of the Southeastern United States. Access to the clinical data was provided by the Rush Alzheimer’s Disease Center (RADC) of the Rush University Medical Center in Chicago, IL. The clinical data provided by RADC comes from the participants in the Religious Orders Study and Rush Memory and Aging Project (ROSMAP). ROSMAP is supported by National Institutes of Health grants P30AG10161, R01AG15819, and R01AG17917, and by the Illinois Department of Public Health. Helpful discussions, advice on statistical analysis, and comments on the manuscript prior to submission by Professors Sue Leurgans at Rush and Ruoqing Zhu at the University of Illinois are gratefully acknowledged.

## References

[1] Anastasio TJ. Editorial: Computational and Experimental Approaches in Multi-target Pharmacology. Front Pharmacol. 2017;8:443.

[2] Schmitt B, Bernhardt T, Moeller H-J, Heuser I, Frölich L. Combination therapy in Alzheimer’s disease. CNS drugs. 2004;18:827–44.

[3] Bajda M, Guzior N, Ignasik M, Malawska B. Multi-target-directed ligands in Alzheimer's disease treatment. Curr Med Chem. 2011;18:4949–75.

[4] Haber M, Abdel Baki SG, Grin’kina NM, Irizarry R, Ershova A, Orsi S, et al. Minocycline plus N-acetylcysteine synergize to modulate inflammation and prevent cognitive and memory deficits in a rat model of mild traumatic brain injury. Exp Neurol. 2013;249:169–77.

[5] Tsoi KK, Chan JY, Leung NW, Hirai HW, Wong SY, Kwok TC. Combination Therapy Showed Limited Superiority Over Monotherapy for Alzheimer Disease: A Meta-analysis of 14 Randomized Trials. J Am Med Dir Assoc. 2016;17:863 e1-8.

[6] Wenzel TJ, Klegeris A. Novel multi-target directed ligand-based strategies for reducing neuroinflammation in Alzheimer’s disease. Life Sci. 2018.

[7] Bennett DA, Buchman AS, Boyle PA, Barnes LL, Wilson RS, Schneider JA. Religious Orders Study and Rush Memory and Aging Project. Journal of Alzheimer’s Disease. 2018:1–28.

[8] Cai Z, Hussain MD, Yan LJ. Microglia, neuroinflammation, and beta-amyloid protein in Alzheimer’s disease. Int J Neurosci. 2014;124:307–21.

[9] Regen F, Hellmann-Regen J, Costantini E, Reale M. Neuroinflammation and Alzheimer’s Disease: Implications for Microglial Activation. Curr Alzheimer Res. 2017;14:1140–8.

[10] Sochocka M, Zwolinska K, Leszek J. The Infectious Etiology of Alzheimer’s Disease. Curr Neuropharmacol. 2017;15:996–1009.

[11] Balducci C, Forloni G. Novel targets in Alzheimer’s disease: A special focus on microglia. Pharmacol Res. 2018;130:402–13.

[12] Kokiko-Cochran ON, Godbout JP. The Inflammatory Continuum of Traumatic Brain Injury and Alzheimer’s Disease. Front Immunol. 2018;9:672.

[13] Labzin LI, Heneka MT, Latz E. Innate Immunity and Neurodegeneration. Annu Rev Med. 2018;69:437–49.

[14] Nakamura Y. Regulating factors for microglial activation. Biol Pharm Bull. 2002;25:945–53.

[15] Chhor V, Le Charpentier T, Lebon S, Ore MV, Celador IL, Josserand J, et al. Characterization of phenotype markers and neuronotoxic potential of polarised primary microglia in vitro. Brain Behav Immun. 2013;32:70–85.

[16] Jiang J, Dingledine R. Prostaglandin receptor EP2 in the crosshairs of anti-inflammation, anti-cancer, and neuroprotection. Trends Pharmacol Sci. 2013;34:413–23.

[17] Heneka MT, Carson MJ, El Khoury J, Landreth GE, Brosseron F, Feinstein DL, et al. Neuroinflammation in Alzheimer’s disease. Lancet Neurol. 2015;14:388–405.

[18] Malm TM, Jay TR, Landreth GE. The evolving biology of microglia in Alzheimer’s disease. Neurotherapeutics. 2015;12:81–93.

[19] Kaminska B, Mota M, Pizzi M. Signal transduction and epigenetic mechanisms in the control of microglia activation during neuroinflammation. Biochim Biophys Acta. 2016;1862:339–51.

[20] Anastasio TJ. Temporal-logic analysis of microglial phenotypic conversion with exposure to amyloid-beta. Mol Biosyst. 2014.

[21] Anastasio TJ. Computational identification of potential multi-drug combinations for reduction of microglial inflammation in Alzheimer disease. Front Pharmacol. 2015;6:116.

[22] Pineda FJ. Generalization of back-propagation to recurrent neural networks. Physical Review Letters. 1987;59:2229–32.

[23] Pineda FJ. Recurrent backpropagation and the dynamical approach to adaptive nerual computation. Neural Comput. 1989;1:161–72.

[24] Perrone MP, Cooper LN. When networks disagree: Ensemble methods for hybrid neural networks. How We Learn; How We Remember: Toward An Understanding Of Brain And Neural Systems: Selected Papers of Leon N Cooper: World Scientific; 1995. p. 342–58.

[25] Butovsky O, Talpalar AE, Ben-Yaakov K, Schwartz M. Activation of microglia by aggregated beta-amyloid or lipopolysaccharide impairs MHC-II expression and renders them cytotoxic whereas IFN-gamma and IL-4 render them protective. Mol Cell Neurosci. 2005;29:381–93.

[26] Piazza A, Lynch MA. Neuroinflammatory changes increase the impact of stressors on neuronal function. Biochem Soc Trans. 2009;37:303–7.

[27] Grinberg YY, Dibbern ME, Levasseur VA, Kraig RP. Insulin-like growth factor-1 abrogates microglial oxidative stress and TNF-alpha responses to spreading depression. J Neurochem. 2013;126:662–72.

[28] Miklossy J. Chronic inflammation and amyloidogenesis in Alzheimer’s disease -- role of Spirochetes. J Alzheimers Dis. 2008;13:381–91.

[29] Bibi F, Yasir M, Sohrab SS, Azhar EI, Al-Qahtani MH, Abuzenadah AM, et al. Link between chronic bacterial inflammation and Alzheimer disease. CNS Neurol Disord Drug Targets. 2014;13:1140–7.

[30] Bennett DA, Wilson RS, Schneider JA, Evans DA, Beckett LA, Aggarwal NT, et al. Natural history of mild cognitive impairment in older persons. Neurology. 2002;59:198–205.

[31] Bennett DA, Schneider JA, Arvanitakis Z, Kelly JF, Aggarwal NT, Shah RC, et al. Neuropathology of older persons without cognitive impairment from two community-based studies. Neurology. 2006;66:1837–44.

[32] Guo Z, Fratiglioni L, Zhu L, Fastbom J, Winblad B, Viitanen M. Occurrence and progression of dementia in a community population aged 75 years and older: relationship of antihypertensive medication use. Arch Neurol. 1999;56:991–6.

[33] Rohde D, Hickey A, Williams D, Bennett K. Cognitive impairment and cardiovascular medication use: Results from wave 1 of The Irish Longitudinal Study on Ageing. Cardiovasc Ther. 2017;35.

[34] Forette F, Seux ML, Staessen JA, Thijs L, Babarskiene MR, Babeanu S, et al. The prevention of dementia with antihypertensive treatment: new evidence from the Systolic Hypertension in Europe (Syst-Eur) study. Arch Intern Med. 2002;162:2046–52.

[35] Hajjar I, Catoe H, Sixta S, Boland R, Johnson D, Hirth V, et al. Cross-sectional and longitudinal association between antihypertensive medications and cognitive impairment in an elderly population. J Gerontol A Biol Sci Med Sci. 2005;60:67–73.

[36] Khachaturian AS, Zandi PP, Lyketsos CG, Hayden KM, Skoog I, Norton MC, et al. Antihypertensive medication use and incident Alzheimer disease: the Cache County Study. Arch Neurol. 2006;63:686–92.

[37] Gelber RP, Ross GW, Petrovitch H, Masaki KH, Launer LJ, White LR. Antihypertensive medication use and risk of cognitive impairment: the Honolulu-Asia Aging Study. Neurology. 2013;81:888–95.

[38] Soto ME, van Kan GA, Nourhashemi F, Gillette-Guyonnet S, Cesari M, Cantet C, et al. Angiotensin-converting enzyme inhibitors and Alzheimer’s disease progression in older adults: results from the Reseau sur la Maladie d’Alzheimer Francais cohort. J Am Geriatr Soc. 2013;61:1482–8.

[39] Yasar S, Xia J, Yao W, Furberg CD, Xue QL, Mercado CI, et al. Antihypertensive drugs decrease risk of Alzheimer disease: Ginkgo Evaluation of Memory Study. Neurology. 2013;81:896–903.

[40] Anekonda TS, Quinn JF. Calcium channel blocking as a therapeutic strategy for Alzheimer’s disease: the case for isradipine. Biochim Biophys Acta. 2011;1812:1584–90.

[41] Nimmrich V, Eckert A. Calcium channel blockers and dementia. Br J Pharmacol. 2013;169:1203–10.

[42] Czirr E, Weggen S. Gamma-secretase modulation with Abeta42-lowering nonsteroidal anti-inflammatory drugs and derived compounds. Neurodegener Dis. 2006;3:298–304.

[43] Imbimbo BP. Therapeutic potential of gamma-secretase inhibitors and modulators. Curr Top Med Chem. 2008;8:54–61.

[44] Imbimbo BP, Giardina GA. Gamma-secretase inhibitors and modulators for the treatment of Alzheimer’s disease: disappointments and hopes. Curr Top Med Chem. 2011;11:1555–70.

[45] Ettcheto M, Sanchez-Lopez E, Pons L, Busquets O, Olloquequi J, Beas-Zarate C, et al. Dexibuprofen prevents neurodegeneration and cognitive decline in APPswe/PS1dE9 through multiple signaling pathways. Redox Biol. 2017;13:345–52.

